# Identification of Salivary Gland Escape Barriers to Western Equine Encephalitis Virus in the Natural Vector, *Culex tarsalis*

**DOI:** 10.1101/2022.01.11.475797

**Authors:** Charles B. Stauft, Aaron T. Phillips, Tony T. Wang, Kenneth E. Olson

## Abstract

Herein we describe a previously uninvestigated salivary gland escape barrier (SEB) in *Culex tarsalis* mosquitoes infected with two different strains of Western equine encephalitis virus (WEEV). The WEEV strains were originally isolated either from mosquitoes (IMP181) or a human patient (McMillan). Both IMP181 and McMillan viruses were fully able to infect the salivary glands of *Culex tarsalis* after intrathoracic injection as determined by expression of mCherry fluorescent protein. IMP181, however, was better adapted to transmission as measured by virus titer in saliva as well as transmission rates in infected mosquitoes. We used chimeric recombinant WEEV strains to show that inclusion of IMP181-derived structural genes partially circumvents the SEB.

**Author Statement:** During the first half of the previous century, WEEV was responsible for large outbreaks throughout the northern United States and Canada that caused severe disease in horses and people. Over the past 60 years, cases of WEEV have mysteriously faded and the pathogen is rarely encountered in the clinic today. Salivary gland escape barriers (SEB) are a relatively neglected field of study in arbovirology, and this study provides a valuable contribution to the field by describing a SEB found in otherwise vector competent *Culex tarsalis* mosquitoes. Although midgut barriers are well studied, less is known about barriers to transmission in the salivary glands. Although salivary gland infection occurs at a high rate following direct injection of virus into the hemocoel, we noticed that only ∼20-30% of infected mosquitoes transmit detectable infectious virus in their saliva. Additionally, although the more pathogenic patient-derived McMillan strain of WEEV infected salivary glands at a similar rate, its transmission was more severely restricted than the mosquito-derived but less pathogenic Imperial 181 strain. We were able to trace determinants of viral transmission to the 6K/E1 region of the gene encoding the viral structural polyprotein. WEEV is a valuable research model for the closely related Eastern equine encephalitis virus and Venezuelan equine encephalitis virus we believe that our findings are applicable to other members of *Togaviridae*.

## Introduction

Western equine encephalitis virus (WEEV) is maintained in an enzootic transmission cycle between *Culex tarsalis* and passerine bird species. Except for a few recorded incidences of viremia in equine species (1–3), infection of humans or horses is considered a dead-end for viral propagation (1). However, WEEV transmission to equine or human hosts has been associated with severe disease outbreaks, and human survivors of encephalitic infection may experience long-term neurological sequelae (4–7). More recently, WEEV McMillan has been used to model viral-induced parkinsonism in a mouse model of disease (8). The molecular determinants of WEEV McMillan neurovirulence in mice have been mapped to the structural protein E2 (9).

In the mosquito, natural infection occurs *per os* through the ingestion of a blood-meal containing infectious virus. The virus is able to infect the midgut of the mosquito, replicate, escape the midgut into the hemocoel, and then infect the salivary glands for a second round of replication followed by release into the salivary gland lumen (10). Midgut entry and escape barriers have previously been observed in WEEV infection of *Cx. tarsalis* (11,12) and a salivary gland infection barrier has also been described (11). These previous studies have stopped short of describing barriers to secretion of infectious virus particles into the saliva. High concentrations of alphaviruses can be detected in mosquito saliva. Depending on the alphavirus and vector species combination, concentrations from 1,000 to 100,000 mouse LD_50_ per mosquito were detected (13–17). A more recent study detected up to 3.6 × 10^7^ PFU per mosquito of VEEV isolated from extracted mosquito saliva (18). Saliva titers of a highly related alphavirus, Highlands J virus, was reported to be 3.89 × 10^3^ PFU/ saliva sample (19). However, the mechanisms of barriers to alphavirus transmission at the salivary gland level remain incompletely understood.

*Cx. tarsalis*, the principal vector for WEEV, is refractory to midgut infection by McMillan (9,20) and permissive for the Imperial 181 (IMP181) strain of WEEV (9). The molecular determinants for IMP181, in terms of mosquito midgut infectivity, exist in the same region of E2 as was identified for neurovirulence in mice. Although the mosquito infectivity and neurovirulence in mice have been reported for McMillan and IMP181 isolates in these earlier studies, there has not been an investigation of salivary gland barriers to transmission. In this report, we describe the existence of a salivary gland escape barrier for WEEV in *Cx. tarsalis* and provide evidence for the role of the structural proteins in negotiating this barrier.

## Materials and methods

### Virus construction and growth curves

The McMillan infectious clone was originally assembled by Dr. Thomas Welte at Colorado State University (CSU). Imperial 181 was isolated in 2005 from a *Culex tarsalis* caught in Imperial County, CA and used to construct an infectious clone which was obtained from Dr. Aaron Brault at the Centers for Disease Control and Prevention.

Reporter viruses were generated to facilitate screening of infected mosquitoes. A reporter gene encoding mCherry was inserted into the multiple cloning site (MCS) of the 5’dsWEEV.McM plasmid to make an alphavirus expression system. The gene encoding mCherry fluorescent protein was also introduced into the MCS downstream of the native 26S sub-genomic promoter of the 5’dsWEEV.IMP181 plasmid. The primers used to amplify mCherry gene inserts for cloning into pMcM-mCherry were 5’-AAAACCGCGGATGGTGAGCAAGGG and AAAACCTGCAGGTTACTTGTACAGCTCG-3’. The primers used to amplify mCherry gene inserts for cloning into pIMP181-mCherry were 5’-AAACCGCGGATGGTGAGCAAGGG-3’ and 5’-AAACCCGGGTTACTTGTACAGCTCG-3’.

Once amplified and sequenced, plasmids were linearized by incubation overnight with MfeI (McMillan; New England Biolabs) or NotI (IMP181; New England Biolabs) at 37°C. Otherwise, generation of virus from the infectious clones was conducted as reported previously (24). Infectious clones of the chimeric viruses were constructed previously (9) and are composed of reciprocal crosses of C/E3/E2 or 6K/E1 regions from McMillan and IMP181 (Fig 1A).

**Figure 1:**
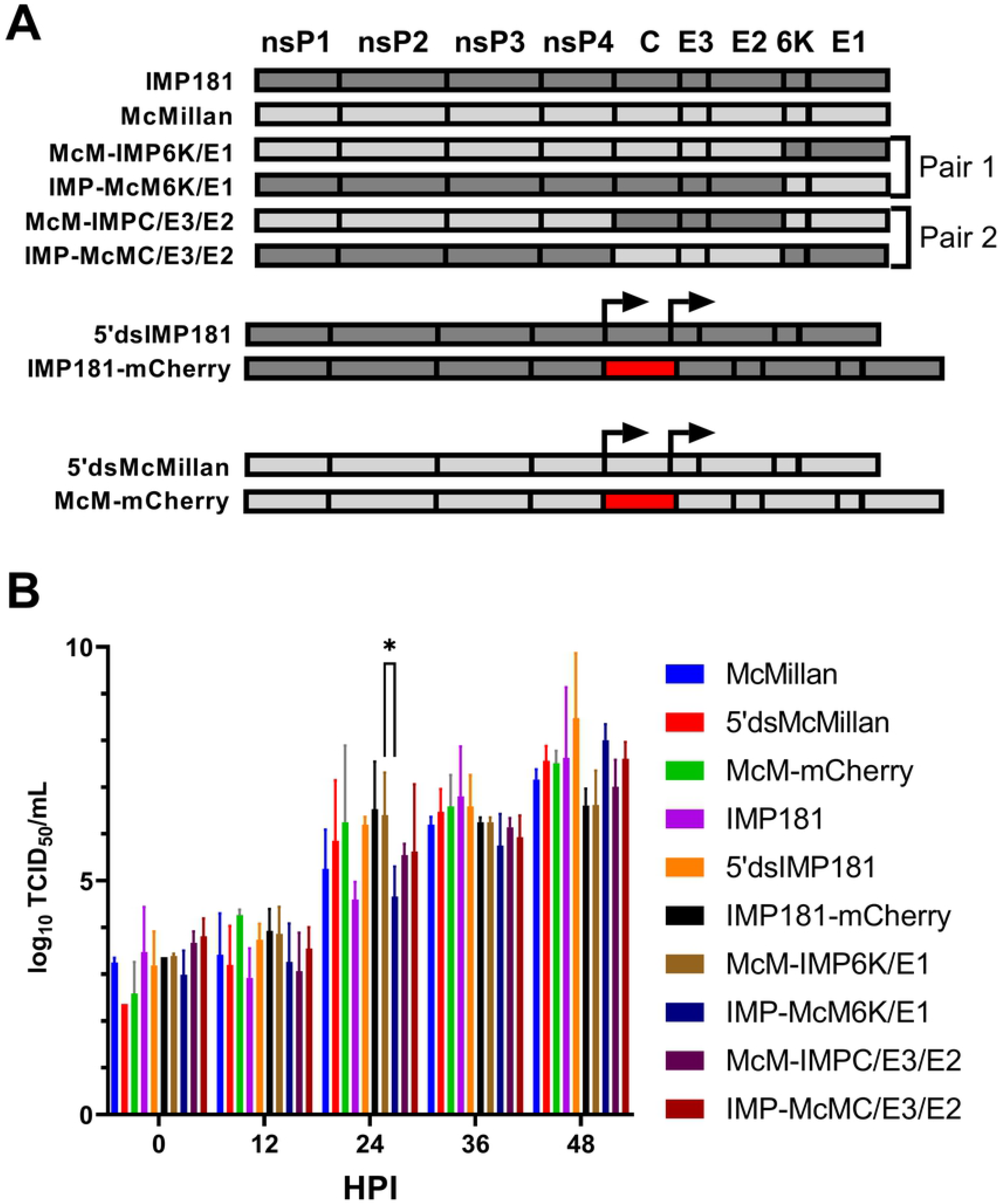
Illustration of WEEV constructs and growth kinetics *in vitro*. A panel of recombinant WEEV constructs (A) were used to study the salivary gland escape barrier in *Culex tarsalis*. The viability of each recombinant virus was assessed using growth kinetics in C6/36 (B) cells. Arrows indicate the native and duplicated sub-genomic promoters. HPI: Hours post-infection.

Growth curves of the WEEV strains were conducted in C6/36 cells (ATCC) grown at 28°C, 5% CO_2_ in MEM (Gibco) supplemented with 10% FBS (Gibco). Confluent monolayers of each cell type were infected with a MOI of 0.01 for each virus and aliquots of supernatant taken every 12 hours for 48 or 60 hours. WEEV was quantified using replicate ten-fold serial dilutions in 90 μL MEM 7% FBS followed by addition of Vero cell (ATCC) suspensions and calculation of TCID_50_/mL using the method of Reed and Muench (21).

### Mosquito infection

*Cx. tarsalis* (Bakersfield, CA) mosquitoes were initially raised in the Arthropod-Borne and Infectious Diseases Laboratory (AIDL), Colorado State University insectaries at 25°C, 80% humidity with a 16:8 light:dark cycle prior to transfer to BSL-3 for WEEV infection at 1 week post-emergence. In BSL-3, the mosquitoes were incubated at 28°C and 80.6% humidity in a Caron 6030 environmental growth chamber. *Cx. tarsalis* were IT injected with 500, 250, or 125 TCID_50_ of McMillan (n=89), McM-mCherry (n=73), IMP181 (n=83), IMP181-mCherry (n=73), McM-IMP6K/E1 (n=78), IMP-McM6K/E1 (n=96), McM-IMPCE3E2 (n=96), or IMP-McMCE3E2 (n=96). At 7 and 14 days post infection mosquitoes were induced to secrete saliva by inserting the proboscis into a capillary tube containing 5 μL of 50% FBS:glycerol solution (15,18).

Salivation proceeded for 30-45 minutes and the salivary glands were dissected for virus detection. The proportion of *Cx. tarsalis* transmitting virus was determined as the number of saliva samples positive for infectious virus divided by the total number of saliva samples. Infected salivary glands were identified by fluorescence microscopy or immune-fluorescence assay (IFA) using an anti-SINV E1 antibody (30.11a). Any mosquitoes found to be negative for salivary gland infection were excluded from further analysis.

Negative controls consisted of saliva collected from non-infected mosquitoes and diluted in MEM with 7% FBS. Positive controls included known doses of McMillan (5 × 10^6^ PFU) strain WEEV added to saliva collected from non-infected mosquitoes either in the capillary tube or after dilution in medium.

### Statistics

At each time point, virus titers from end-point assays were compared using a Student’s T-test for data with normal distributions and Satterthwaite T-test for data lacking a normal distribution. Infectious virus titers in saliva were log transformed and subjected to statistical analysis using Student’s t-test with a post hoc Bonferroni adjustment. The Bonferroni adjustment was used to counteract the loss of statistical power inherent in making multiple statistical comparisons and resulted in an alpha (and a significant p-value) of 0.01. Graphs were assembled using GraphPad Prism (La Jolla, California).

Rates of transmission were compared between each set of related groups (14dpi versus 7dpi, McMillan versus IMP181, IMP181-mCherry versus IMP181, McM-mCherry versus McMillan, mosquitoes inoculated with 500 TCID_50_ versus 250 TCID_50_ and 125 TCID_50_ infected mosquitoes) using confidence intervals of binomial proportions and Z-tests using statistical analysis software (SAS).

## RESULTS

### Virus construction and growth in cell culture

To develop an effective model to study the interactions of WEEV with its natural vector, double sub-genomic recombinant viruses expressing mCherry were constructed. The two principal strains of WEEV used in this study were isolated originally either from a human (McMillan) or *Cx. tarsalis* (IMP181) and propagated in mouse brains or Vero cells respectively an unknown number of times (4,22,23) before being used to construct infectious clones (24,25). McMillan and IMP181 have 99.7% identity at the nucleotide sequence level and significant amino acid sequence identity. McMillan and IMP181 differ greatly in their ability to cause disease in an outbred CD-1 mouse model of infection. While both strains are neuroinvasive, IMP181 caused no mortality in mice after intranasal or subcutaneous infection but McMillan caused high mortality within 5 days (26). Enzootic strains such as IMP181 are associated with a lack of neurovirulence in mice (4). The McMillan strain was originally isolated in 1941 and has been used in numerous studies of WEEV (4,26–28). The growth kinetics for wildtype and infectious clone derived IMP181 and McMillan as well as chimeric recombinants and mCherry expressing constructs were compared in mosquito derived C6/36 cells (Fig 1B). All viruses demonstrated similar growth kinetics with a logarithmic increase in titers through the 48-hour time point (Fig 1B). Additionally, growth kinetics of McM-mCherry and IMP181-mCherry were compared to wild-type McMillan and IMP181 viruses. Peak titers were not significantly reduced in recombinant virus strains in C6/36 cells despite the addition of a second sub-genomic promoter and the gene encoding mCherry fluorescent protein.

### Salivary gland infection

Adult, female, *Cx. tarsalis* mosquitoes were intrathoracically injected to avoid complications with the McMillan strain of WEEV, which is unable to efficiently infect the midguts of *Cx. tarsalis* (9,20). To establish parity between the wildtype strains and reporter-expressing recombinant viruses (McM-mCherry and IMP181-mCherry), we confirmed that the salivary gland infection rate for McMillan and IMP181 compared to McM-mCherry or IMP181-mCherry viruses were not significantly different 7 days following injection with 500 TCID_50_ (n=73-96, Fig 2). Additionally, McM-mCherry and IMP181-mCherry viruses displayed similar patterns of mCherry expression with uniform expression in the distal and proximal lateral lobes and the medial lobe of each set of salivary glands examined at 7 DPI (Fig 2B and 2C). In addition to mCherry expression, anti-SINV E1 antibody (30.11a) also revealed an almost complete infection of the salivary glands by both strains of virus at 7 DPI (Fig 2F and 2H). The 30.11a antibody also recognizes McMillan and IMP181 E1 due to the recombinant nature of WEEV. In salivary glands that were incompletely infected, the distal tips of the lateral lobes exhibited a lack of fluorescence. Prominent tissues expressing viral antigen included acinar cells and duct tissue connecting the glands to the proboscis.

**Figure 2:**
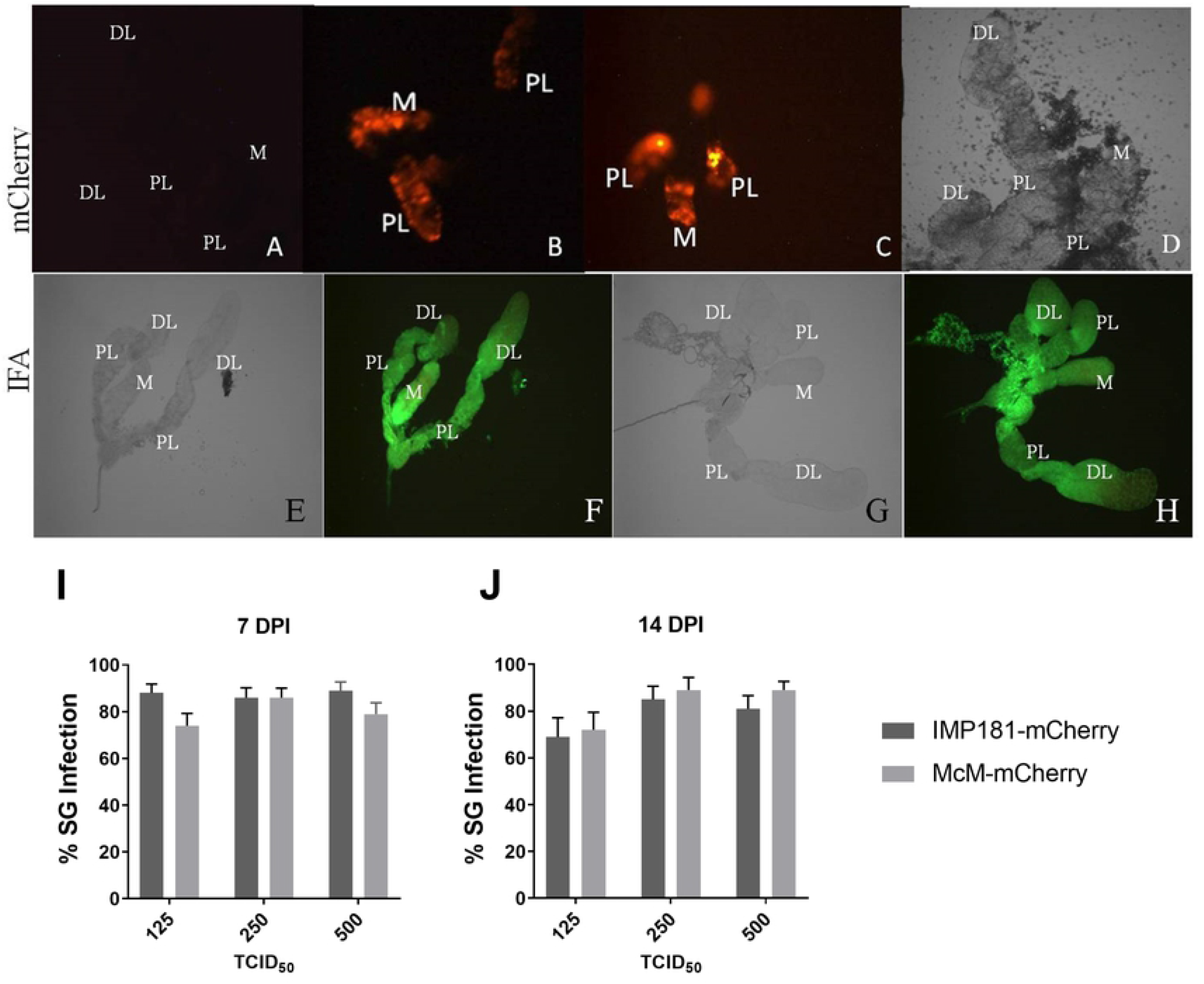
Epifluorescent imaging of *Cx. tarsalis* salivary glands and rates of salivary gland infection. Salivary glands were compared to negative control epifluorescent (A) and light (D) images of non-infected *Cx. tarsalis* salivary glands at 20x magnification. *Cx. tarsalis* salivary glands 7 days post intrathoracic injection with 500 TCID_50_ McM-mCherry (B) and IMP181-mCherry (C). IFA of *Cx. tarsalis* salivary glands. *Cx. tarsalis* salivary glands stained immunohistochemically for WEEV 7 days post intrathoracic injection with 500 TCID_50_ McMillan (F) and IMP181 (H) along with light microscopy images of the same salivary glands (E and G). Salivary gland infection rates were calculated using the number of fluorescent mosquitoes compared to total injected mosquitoes at 7 (I) and 14 (J) DPI. DL, distal-lateral lobe; M, medial lobe; PL, proximal-lateral lobe. DPI: Days post-infection.

Epifluorescent imaging of salivary glands was conducted to determine the presence or absence of a salivary gland infection barrier (SIB) in *Cx. tarsalis*. Salivary gland infection by McM-mCherry or IMP181-mCherry would be marked by the presence or absence of mCherry expression. There were no visible differences in the pattern of salivary gland infection or frequency of infected salivary glands between IMP181-mCherry and McM-mCherry (Fig 2). Salivary gland infection rates associated with each virus were not significantly different and ranged from 74% to 89%, indicating the absence of a significant SIB. The rates of salivary gland infection were not significantly different between different doses (125, 250, and 500 TCID_50_) of McM-mCherry and IMP181-mCherry at 7 days (Fig 2I) or 14 days post infection (Fig 2J).

Reporter expressing recombinant viruses (IMP181-mCherry and McM-mCherry) were used to assess the impact of varied dosage and duration of infection on transmission rates as measured by the percentage of saliva containing detectable infectious virus. Rates of transmission were not found to vary significantly within groups of *Cx. tarsalis* injected with different doses of IMP181-mCherry. However, McM-mCherry infected mosquitoes showed a significant increase in transmission rate from doses of 250 to 500 TCID50 at 7 and 14 days post injection (Fig 3A and 3B). Transmission rates were significantly greater for IMP181-mCherry than McM-mCherry (Fig 3A) at 7 days post injection with 125, 250, and 500 TCID_50_ (Fig 3). After 14 DPI, transmission rates were found to be higher in IMP181-mCherry compared to McM-mCherry infected mosquitoes at the lowest dose of 125 TCID50, however, transmission rates were similar at the 250 and 500 TCID50 doses. The transmission rate was found to be dose dependent with McM-mCherry, however, no dose-dependence was observed in the dose range attempted for IMP181-mCherry infections (Fig 3A and 3B).

**Figure 3:**
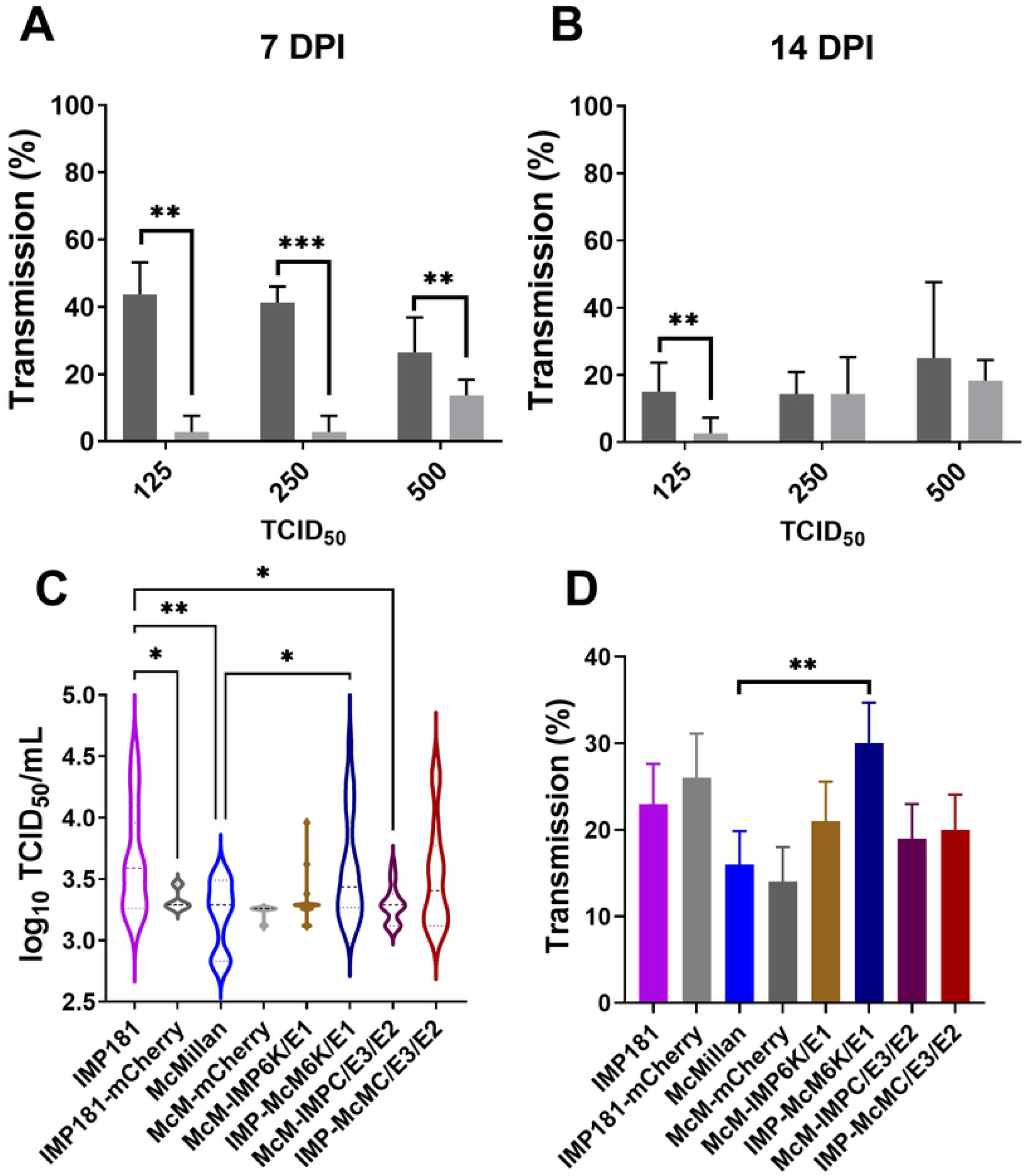
Salivary gland infection rates and infectious virus concentration in saliva. Salivary gland infection rates at 7 (A) and 14 (B) days were compared between IMP181-mCherry (dark grey) and McMillan (light grey) infected mosquitoes injected with 125, 250, or 500 TCID_50_. Infectious viral titers were also compared between IMP181 and McMillan derived viruses in saliva collected at 7 days post-injection with 500 TCID50 with IMP181 6K/E1 genes associated with an increase in expectorated virus (C). Transmission rates were also assessed for IMP181 and McMillan derived viruses (D).

Despite similarity in rates of salivary gland infection, significant differences were observed in the concentration of infectious WEEV recovered in mosquito saliva (Fig 3). IMP181 infected mosquitoes secreted significantly more virus in their saliva compared to McMillan injected mosquitoes at 7 DPI (p=0.0011; Student’s T test). Following up on this difference, saliva virus titers were measured for chimeric McM-IMP181 recombinant viruses. IMP-McM6K/E1 (p=0.0010) and IMP-McMC/E3/E2 (p=0.0066) had higher infectious virus titers in saliva compared to wild-type McMillan at 7 days post injection. IMP-McM6K/E1 and IMP-McMC/E3/E2 also demonstrated significantly different saliva titers when compared to IMP181 (p=0.2602 IMP-McM6K/E1; 0.1706 IMP-McMC/E3/E2; Fig 3C). Despite differences in the concentration of expectorated virus, transmission rates were broadly similar between groups **(Figure 3D)**. Although approaching significance in a one-tailed test comparing binomials (p=0.0504), IMP181 transmission rates were not found to be higher than those in McMillan infected mosquitoes. However, a significant difference was observed in IMP-McM6K/E1 infected mosquitoes compared to the McMillan group (Fig 3D). In these chimeric recombinant viruses, increases in salivary gland titers associated with the IMP181 structural genes for 6K and E1. We did not observe significantly different concentrations of infectious virus in the saliva of McM-IMPC/E3/E2 or IMP-McM6K/E1, which possesses the nonstructural and 6K/E1 regions of McMillan. A protein BLAST comparing the McMillan (ACT75276) and IMP181 (ACT75278) structural polyproteins reveals 32/1236 amino acid mismatches. Of these mismatches 22 are in the C/E3/E2 region and 10 are in 6K/E1. In future studies, further subcloning and site-directed mutagenesis may enable the location of key mutations involved in egress from infected salivary glands.

## Conclusions

A salivary gland infection barrier (SIB) was not detected with the *Culex tarsalis* CA strain for McMillan or IMP181 after intrathoracic injection as salivary gland infection was not shown to vary by dose, time, or strain of virus in this study. This observation agrees with previous work **(11,12)**. Both McM-mCherry and IMP181-mCherry could infect *Cx. tarsalis* salivary glands and express a fluorescent reporter that allowed for clear demarcation of infected versus uninfected tissue that was comparable to IFA. Dose dependence of transmission suggested a relationship between virus titer in the hemocoel to the expectoration of virus into the saliva of infected mosquitoes. Time after infection (7 or 14 days post intrathoracic injection) was not shown to significantly affect salivary gland infection, transmission rate in saliva, or amount of WEEV expectorated.

Previous work **(29)** identified a SIB in a different strain of *Cx. tarsalis*. Our work identified a salivary gland escape barrier to WEEV in the CA strain of *Cx. tarsalis*. An escape barrier was likely responsible for the lack of virus secreted in the saliva of infected mosquitoes as rates of transmission varied between 14-30% while salivary gland infection averaged around 74-89% for both strains of WEEV. The amount of virus detected in saliva was significantly greater for IMP181, clone 40, and clone 42 virus infected mosquitoes. IMP181-derived structural genes appeared to have an effect with regards to the phenotype of increased expectoration of virus in saliva as transmission of McMillan was enhanced by the structural genes of IMP181. Both halves of the structural gene encoding region were equally important to replication or egress in the salivary gland tissue of *Culex tarsalis*. In agreement with prior work **(9)**, the IMP181 strain appears to be more highly adapted to replication in the mosquito vector than the highly passaged clinical isolate strain McMillan. The McMillan strain likely has adapted to vertebrate cells after lengthy passaging under laboratory conditions and therefore is a useful counterpoint for studying wild-type isolates like IMP181 which have maintained the ability to infect and be transmitted by competent vectors. IMP181 and McMillan form a dyad in the spectrum between vector infectivity and vertebrate host virulence that have been used in past studies to yield useful conclusions.

IFA revealed the presence of WEEV E1 along the walls of the salivary gland duct for McMillan and IMP181 infected mosquitoes. As IFA is not quantitative, it is not possible to separate the roles of budding and encapsidation in determining the amount of WEEV egress in this study. Infectious virus titer in the saliva did not increase significantly with dose of IMP181-mCherry or McM-mCherry injected into the mosquito. This indicated that the limiting step for WEEV egress occurred within the salivary gland as the amount of WEEV in the hemocoel did not alter the amount of expectorated virus. Future studies using IFA with antibodies specific for different WEEV structural proteins and confocal microscopy could contribute enhanced visualization to the study of viral escape from infected salivary glands.

Transmission rate was shown to be related to inoculation dosage in McM-mCherry as rates were significantly higher in 500 TCID_50_ injected mosquitoes. Dose dependence was not observed with the transmission rate of IMP181-mCherry. Homogenous presentation of IMP181-mCherry transmission was possibly due to injection doses being higher than the minimum threshold for salivary gland infection, replication, or escape. Also, *Cx. tarsalis* has been shown to be more susceptible to IMP181 compared to McMillan in terms of midgut infection (9). The 5’ (AvrII to KpnI) or 3’ (KpnI to terminus) sections of the structural polyprotein of IMP181 previously rescued mosquito infection with either McMillan or IMP181 backgrounds (9). The same two regions have been shown in this study to be involved with salivary gland escape of WEEV in the saliva. Midgut barriers are encountered much earlier and so overshadow the role of the salivary glands in arboviral transmission cycles. However, transmission of infectious virus in this study was limited in infected CA strain *Cx. tarsalis* mosquitoes by a salivary gland escape barrier. Additionally, key regions of the WEEV genome were found to be associated with viral evasion of the SEB that are also involved in vertebrate host pathogenesis. As enzootic and epizootic transmission of WEEV in nature continues to wane, additional research is needed to identify key determinants of decline and apply those findings to other arthropod-borne viruses.

## Acknowledgements

We would like to thank Irma Sanchez-Vargas for assistance in dissecting mosquito salivary glands.

